# Accuracy Feedback and Delay Adaptation Effects in Visual and Tactile Duration Reproduction

**DOI:** 10.1101/2023.12.04.569859

**Authors:** Lingyue Chen, Stephanie Brunner, Zhuanghua Shi

**Affiliations:** Ludwig-Maximilians-Universität München; Vocational school for emergency paramedics at the Bavarian Red Cross, Munich, Germany

**Keywords:** Time perception, accuracy feedback, delay adaptation, multisensory and motor integration

## Abstract

Brief actions involve multiple temporal cues that may not always synchronize, and in basic action-effect relationships, the effect is often delayed. How the brain incorporates delays across modalities in a sensorimotor duration reproduction remains unclear. To investigate this, we conducted two experiments on duration reproduction with delayed sensory feedback. Participants reproduced durations in either visual (Experiment 1) or tactile (Experiment 2) modalities. In the adaptation phase, a contingent sensory feedback, either visual or tactile, was delayed by 150 ms in one session and synchronized in the control session, with accuracy feedback provided after each trial. In the testing phase, random action-effect delays (0-150 ms) were introduced and accuracy feedback was removed. The findings revealed that accuracy feedback effectively recalibrated motor time but did not eliminate the delay. Without accuracy feedback, tactile reproduction relied more on the tactile feedback than on motor time, resulting in greater lengthening of motor reproduction compared to the visual feedback. These findings suggest that temporal delay adaptation is influenced by accuracy feedback and sensorimotor integration, with sensorimotor reliability assigning a higher weight to the tactile than the visual modality.

## 1. Introduction

Perceiving time is integral to our everyday activities, not just for lengthy experiences like watching a movie but also for fleeting moments lasting only seconds, especially tied to actions (Buhusi & Meck, 2005; Merchant & Yarrow, 2016). Yet, our subjective time does not always match objective time. Several factors, such as stimulus intensity, motion, emotional states, and voluntary actions, influence how we perceive time (Eagleman, 2008; Johnston et al., 2006; Meck, 1983; Park et al., 2003; Shi et al., 2012; Yarrow et al., 2001). Moreover, time perception varies across different sensory modalities (Issa et al., 2020; Johnston et al., 2006; Ogden et al., 2010; Paton & Buonomano, 2018; Wearden et al., 2006). Research has pinpointed multiple brain regions involved in timing tasks, suggesting that time processing is distributed rather than governed by an amodal ‘inner clock’ (Bueti, 2011; Bueti et al., 2008; Lewis & Miall, 2009). For example, a given duration may seem shorter when experienced visually or tactilely as opposed to auditorily (Jones et al., 2009; Walker & Scott, 1981; Wearden et al., 1998). Additionally, it has been suggested that the timing of actions and motor movements may differ from the timing of perceived sensations (Bueti & Walsh, 2010).

When navigating real environments, people simultaneously process timing cues from both internal motor actions and the external sensory feedback they generate. Integrating multiple sensory and motor temporal cues into a coherent percept is fundamental. When we push a light button, we not just feel the ‘push’ from the hand but also see the light on immediately. Other complex actions, like playing piano, demand adeptly coordinating multiple temporal aspects. Players must generate internal prior to coordinate finger movements and pedal work while processing external timing cues from visually interpreting sheet music, audibly discerning the produced tunes, and sensing the tactile feedback from the keys. These integrated internal and external actions and multisensory cues supply crucial temporal information for a captivating musical performance. Even though timing may vary across sensory modalities and between sensory and motor processes, how do we often experience a coherent time perception when bombarded with various multisensory inputs and sensorimotor actions? A prevailing theory posits that the brain combines all available information to form coherent perception based on their reliabilities, as suggested by Bayesian inference models (Ernst & Banks, 2002; Jazayeri & Shadlen, 2010; Shi, Church, et al., 2013), the idea can be dated back to Helmholtz’s “perception as inference” (Helmholtz, 1867).

Yet, sensory inputs and actions are not always perfectly synchronized. For instance, during a virtual meeting with a shaky internet connection, a person might notice a disconnect between the video and audio or a lag between their speech and the corresponding video and audio feedback. Such temporal misalignments can provoke a sensation of ‘cognitive dissonance’ (Festinger, 1962). To counter this unsettling feeling of ‘cognitive dissonance’, our brain strives to resolve the conflicting information into a coherent perception. Remarkably, adapting to environments with temporal delay, such as during long-distance video conferences, can significantly diminish or even nullify our initial perception of that delay. Research by Cunningham and colleagues (2001) illustrates this adaptation. Participants engaged in a simulated shooting video game with a 235 ms delay in the visual feedback. Initially they performed badly and eventually recalibrated sensorimotor delay and reached comparable performance as the control group at the end of the delay adaptation. However, when the delay was removed, their performance drastically declined to about 52% accuracy, exhibiting a “negative aftereffect”. Similarly, Stetson et al. (2006) found a negative aftereffect in an action-feedback delay task. Participants recalibrated their perception of sensorimotor synchronicity when pressing a button produced a delayed flash. In a subsequent test, after this delay was removed, participants felt that the resultant flash from their action occurred before the action nearly 40% of the time. This illusory inversion of action and effect underscores the adaptability of our sense of timing to various environments.

To investigate duration perception, Ganzenmüller et al. (2012) desynchronizing reproduction action and the action-induced event (the sensory feedback that is directly caused by the motor action) using an adaptation-testing paradigm. They found that an onset delay (between action start and feedback onset) caused participants to overproduce the target duration, while an offset delay caused underproduction, to a lesser extent. Interestingly, participants nearly lengthened the reproduced duration by the amount of the onset delay within the same modality. This lengthening effect persisted as long as the delay was present, showing immediate adaptation. In other words, individuals may focus on comparing the duration of the probe event and the duration of the contingent sensory feedback (i.e., within-modality sensory comparison), disregarding delay and lengthening their motor action, even though instructions emphasized the importance of accurately reproducing motor time. One potential reason for such an effect is that Ganzenmüller et al. (2012) did not provide feedback on the accuracy of each motor time reproduction. Comparing durations within the same modality might be easier than comparing sensory and motor time, leading participants to shift to within-modality sensory comparison rather than sensorimotor comparison. This raises several important yet unanswered questions: Would accuracy feedback recalibrate the internal prior for delayed-output reproduction? After delay recalibration, how does the brain treat the delay in the absence of accuracy feedback? Would the recalibrated motor timing shift back to its original state without the delay? Does sensory modality play a role in the integration of delayed feedback?

To formally interpret these findings and our hypotheses, we frame the problem within a Bayesian sensorimotor integration framework (Ganzenmüller et al., 2012; Körding & Wolpert, 2004; Shi, Church, et al., 2013). This approach posits that the brain optimally combines multiple noisy temporal estimates by weighting them according to their relative reliability (precision), a process equivalent to maximum likelihood estimation (MLE). In duration reproduction with delayed sensory feedback, three temporal components are critical: (1) *motor timing* (M), the perceived duration of one’s own motor actions (pressing and holding the button); (2) *external sensory feedback timing* (F), the perceived duration of the action-induced sensory event (the visual or tactile stimulus that appears during button press); and (3) *delay* (D), the temporal discrepancy introduced between the onset of the motor action and the onset of the sensory feedback. Crucially, when a delay is introduced, the motor action duration exceeds the sensory feedback duration by exactly this delay: *M = F+D*.

According to MLE, the reproduced duration emerges from integrating motor timing and sensory feedback timing, with each weighted by its reliability. The weight assigned to sensory feedback depends on modality-specific precision: more reliable sensory information (e.g., tactile vs. visual) receives greater weight in determining the reproduced duration. This integration process is central to understanding how the brain resolves conflicts between motor and sensory temporal information.

During adaptation, in which accuracy feedback was provided for reproduction performance, participants are expected to adjust their motor output to meet task demands despite the introduced delay. Based on previous temporal recalibration studies (Cunningham et al., 2001; Stetson et al., 2006), we anticipated that accuracy feedback would recalibrate internal prior (the internal duration representation) during the adaptation, and thus adjust motor timing, though we were uncertain about the extent of this recalibration. Specifically, during adaptation, participants must reconcile the conflicts among: the probe duration (P) they aim to reproduce, their motor timing (M), and the contingent sensory feedback (F) that differs from M by the delay (D). We hypothesize that participants would partially integrate the delay, such that the effective discrepancy between probe duration and sensory feedback reflects both a general reproduction bias (Δ) and a weighted contribution of the delay:

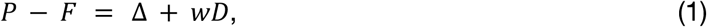

where *w* represents the weight assigned to external sensory feedback timing in determining the motor response. A weight of w=0 indicates complete reliance on internal prior (ignoring any delay in the sensory feedback), while w=1 indicates full trust in contingent sensory feedback (lengthening motor action to compensate for the delay, as observed by Ganzenmüller et al., 2012 without accuracy feedback). By comparing delayed and synchronized conditions, assuming similar general bias Δ, we can estimate the sensory feedback weight during accuracy-feedback-guided recalibration.

In the absence of accuracy feedback during the testing phase, we expected the two phenomena. First, the recalibrated internal prior from the adaptation phase would gradually return to its baseline. Second, and more critically for understanding sensorimotor integration, we expect the MLE framework to govern how motor timing and sensory feedback timing are combined (Ganzenmüller et al., 2012; Körding & Wolpert, 2004; Shi, Church, et al., 2013). Specifically, the perceived duration (D_p_) should emerge as a weighted integration of motor timing (M) and the sensory feedback timing (F):

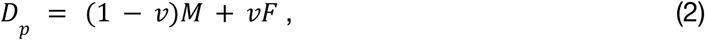

where *v* is the weight assigned to sensory feedback. Participants then compared the perceived duration with the learned prior. Given the fixed relationship between motor timing and sensory feedback timing under delay (*D*), *F = M-D*, Equation (2) can be rewritten as:

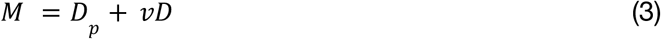

This reformulation reveals that the slope of reproduced motor duration (M) as a function of delay directly reflects *v*. A slope near 1 (*v* ≈ 1) indicates strong reliance on sensory feedback timing, while a slope near 0 (*v* ≈ 0) indicates strong reliance on motor timing. Critically, according to MLE principles, these weights should depend on the relative reliability of each modality’s sensory feedback (Ernst & Banks, 2002; Körding & Wolpert, 2004; Shi, Ganzenmüller, et al., 2013) and modality appropriateness (Welch & Warren, 1980). For example, studies have shown touch had a stronger influence on vision than vision on touch on sequent of visual and tactile events (Bresciani et al., 2006). By varying the delay systematically, we could estimate these modality-specific integration weights.

In summary, we developed three specific hypotheses. First, accuracy feedback during adaptation would recalibrate internal prior to compensate for the introduced delay partially, with the extent of compensation reflected in the weight *w* (Equation 1). Second, in the absence of accuracy feedback during testing, the recalibrated internal prior would gradually decay back to its baseline state. Third, the weight assigned to motor timing versus sensory feedback timing in the MLE integration model (parameter *v* in Equations 2 and 3) would be modality-specific, with tactile feedback receiving a higher weight than visual feedback due to higher sensory reliability. To validate these hypotheses, we conducted two experiments with visual (Experiment 1) and tactile (Experiment 2) feedback modalities. In both experiments, participants reproduced a standard duration and received action-induced feedback (visual or tactile) during reproduction. During adaptation, we provided accuracy feedback and compared delayed (150 ms onset delay) versus synchronized feedback. During testing, we removed the accuracy feedback and systematically varied the onset delay (0-150 ms) to estimate modality-specific integration weights.

## 2. Method

### 2.1. Participants

To avoid potential cross-modal transfer effects, we recruited different groups of participants for the visual (Experiment 1) and tactile (Experiment 2) duration reproduction tasks. Each group initially included 20 participants, but one in each group failed to complete the testing.

Thus, each group ended up with 19 participants (Experiment 1: ages 19-31, mean 24.1, eight females and 11 males; Experiment 2: ages 21-33, mean 25.7, 12 females and seven males). All participants had self-reported normal or corrected-to-normal vision and were naive to the purpose of the experiments. They provided informed consent before the experiment and received 9 Euros per hour for their participation. The experiment was approved by the Ethics Committee.

The sample size was determined based on previous studies. A previous study (Ganzenmüller et al., 2012) found a large effect size for visual delay versus synchronous reproduction feedback (η^2^ =.769, *f* = 1.82). Another study (Tomassini et al., 2011) also found a large effect size for modality differences between visual and tactile durations (η^2^ =. 67, *f* = 1. 42). With a significant level α =. 05 and a statistical power of 95%, our calculations for between-subject comparison showed that at least 15 participants were needed to achieve the desired power. To ensure robust results, we increased the group size to 20 participants.

### 2.2. Apparatus

The experiments took place in a sound-isolated dark cabin. We developed the experimental program using the Psychopy package (Peirce & MacAskill, 2018) in PyCharm. Participants viewed visual stimuli (Gabor patch) on a ViewPixx LCD monitor (VPixx Technologies Inc., Saint-Bruno, Canada) with a refresh rate of 120 Hz, with a consistent viewing distance of 60 cm, maintained by a chin rest, and they provided their behavioral responses using a standard keyboard. Tactile stimuli were presented with an AEC TACTAID VBW32 vibrator attached to the response finger, connected to the computer via an audio amplifier. We placed the tactile stimulation close to the response to enhance the spatial proximity of sensorimotor integration. Participants wore foam earplugs to prevent irrelevant vibration and sound interference and put their arms on a pillow.

### 2.3. Stimulus

Both Experiments 1 and 2 applied the same paradigm but employed stimuli from different sensory modalities. Experiment 1 used a visual Gabor patch (size: 1.7° of visual angle, spatial frequency: 0.08 cycles per degree) on the gray background to reduce afterimage effects. Experiment 2 used tactile vibration (250 Hz) as the primary stimulus for target duration and sensory feedback, while visual fixation served as the trial start signal. In both experiments, the target duration presented for participants to reproduce remains 800 ms per trial. This duration is well-established in temporal reproduction studies (Chen & Shi, 2023; Ganzenmüller et al., 2012; Ren et al., 2021) and provides a sufficient time window to effectively manipulate and analyze the impact of delayed sensory feedback. Furthermore, the same type of stimulus (either visual or tactile) also served as an action-induced sensory feedback when participants began to reproduce the duration.

### 2.4. Procedure

Each experiment consisted of two sessions: a synchronized session and a delayed session. These sessions were separated by a minimum 5-minutes break, and their order was counterbalanced among participants. Each session included an adaptation phase and a testing phase, both involving the same duration reproduction task (Figure 1).

**Figure 1.**
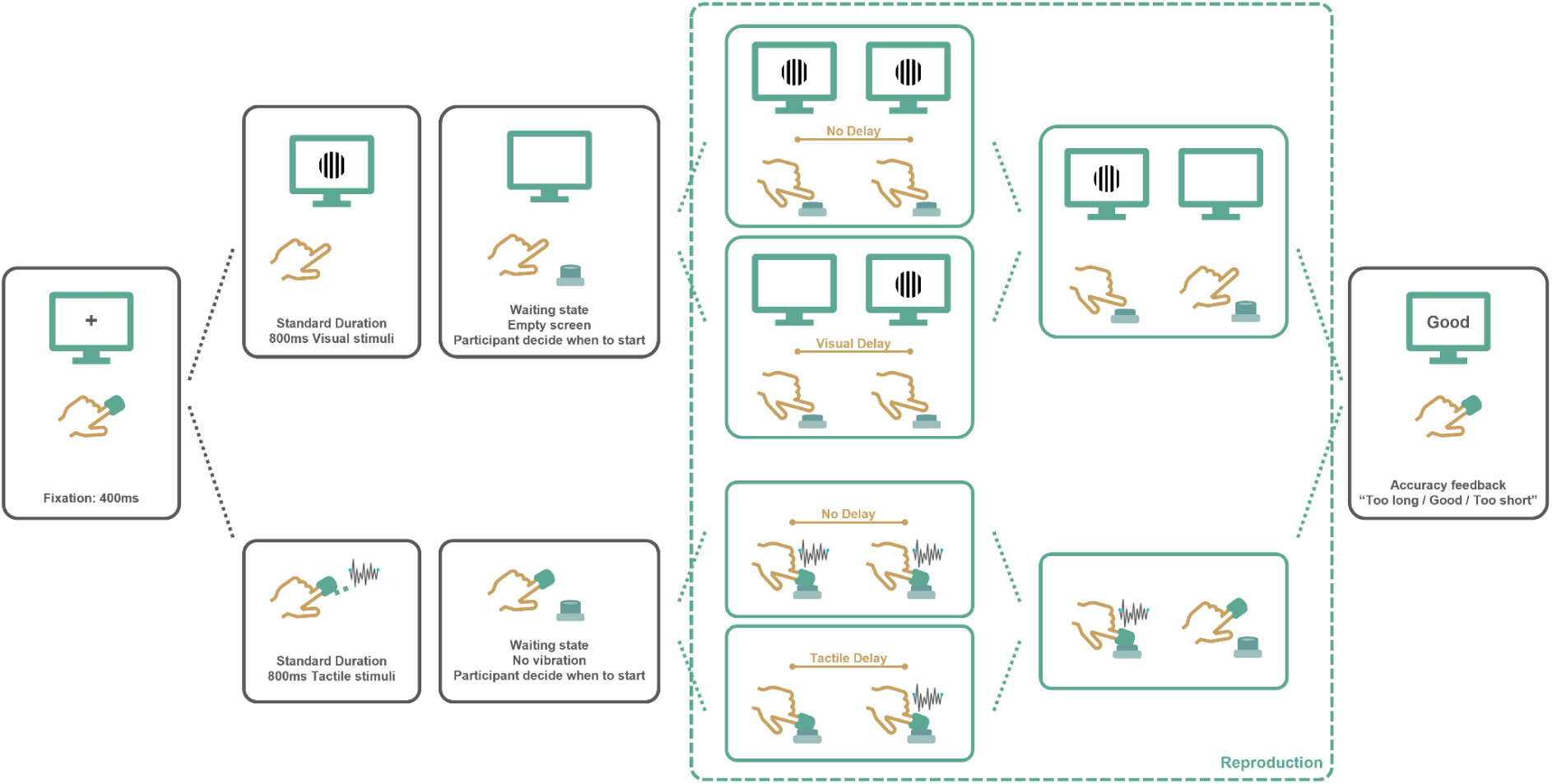
Schematic illustration of experimental procedure. The two experiments shared the same paradigm but differed in modalities through which stimuli were presented (Experiment 1: visual, upper panel; Experiment 2: tactile, bottom panel). In a reproduction task, participants first fixated on a central cross for 400 ms, then a stimulus was presented for 800 ms. After the stimulus disappeared, participants were asked to reproduce the perceived duration as accurately as possible. In both experiments, the action-induced visual or tactile feedback appeared immediately when participants started the reproduction (pressing down the button) in the synchronized session, while an onset delay was introduced in the delayed session. The offset of the action-induced visual or tactile feedback and participants’ reproduction (releasing the button) is always synchronized.

Each trial began with a 400-ms fixation cross, then the first stimulus (a Gabor patch or a vibration) appeared for a fixed duration of 800 ms. After the stimulus disappeared, participants pressed and held the “space” key at their own pace to reproduce the duration as accurately as possible. Upon button press, depending on the session and the phase, the same Gabor patch or vibration appeared immediately or with a delay. The stimulus disappeared as soon as the button was released. During the adaptation phase, the contingent sensory feedback (visual or tactile) appeared immediately in the synchronized session but was delayed 150 ms in the delayed session. In other words, the contingent sensory event was 150 ms shorter than the actual action duration (Figure 2). During the testing phase, however, the contingent sensory events were delayed by a random interval of 0, 50, 100, or 150 ms for both sessions.

**Figure 2.**
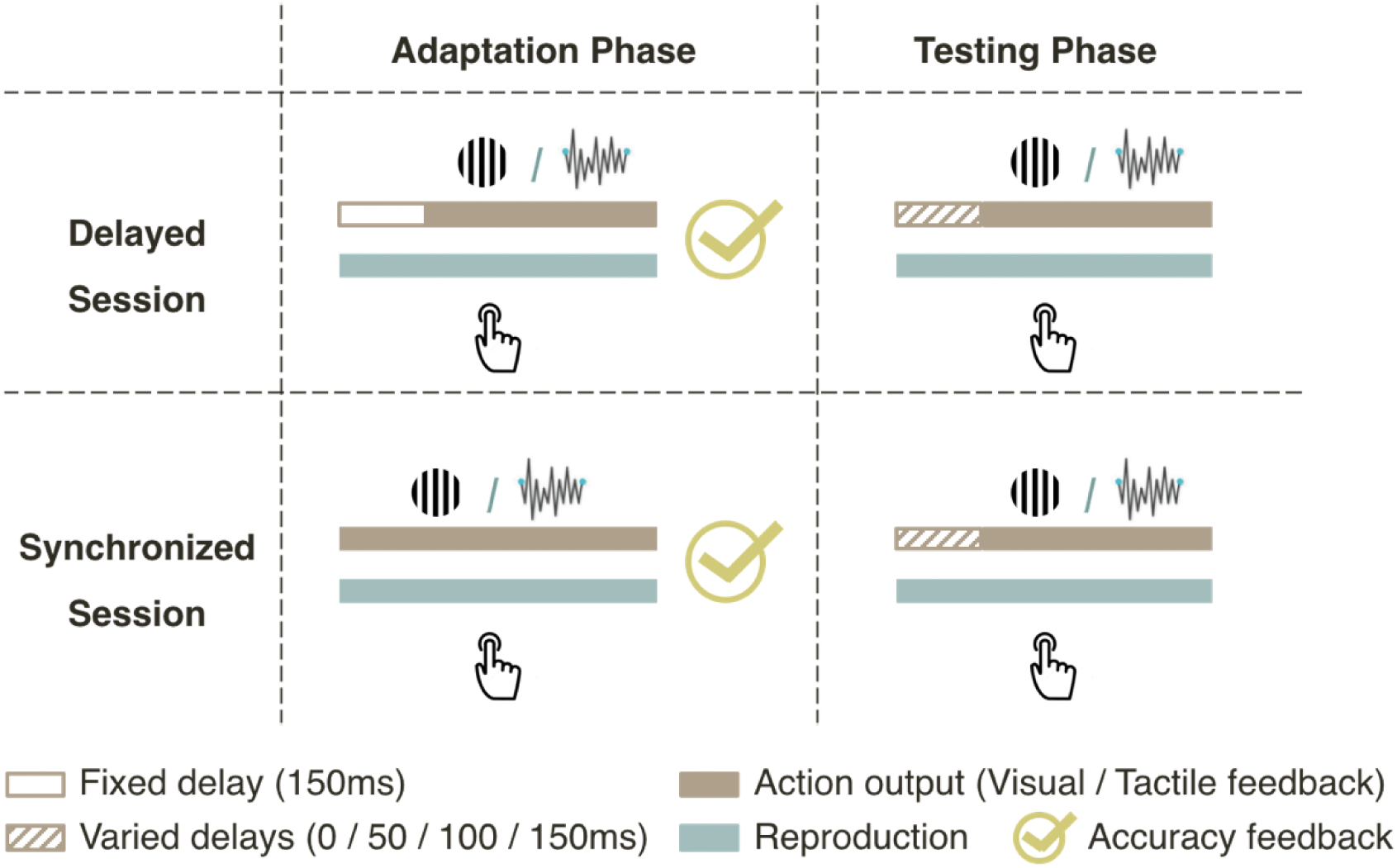
Schematic illustration of the delay manipulation during reproduction. Each experiment consisted of an adaptation phase (left panel), followed by a testing phase (right panel). In the adaptation phase, the contingent sensory (visual / tactile) feedback was presented either with a fixed delay of 150 ms (Delayed session) or synchronized with the onset of the action (Synchronized session). In the testing phase, the delay between action and contingent sensory feedback varied from 0 to 150 ms. Participants received accuracy feedback immediately after the reproduction (denoted by the “tick” symbol) in the adaptation trials while no accuracy feedback was provided after the reproduction in the testing trials.

The adaptation phase included 60 obligatory trials. Participants reproduced these trials and received text feedback on their reproduction accuracy. The adaptation phase ended if 80% of the last ten trials fell within the “good” window. Otherwise, 10 additional trials were added until they reached the 80% criterion. In Experiment 1, 68% of participants reached the goal within 60 trials, and 79% reached the goal within one round of 10 trials. Three participants took more than two rounds to reach the goal. In Experiment 2, all participants reached the goal within the initial 60 trials.

The testing phase began automatically after the adaptation phase. It consisted of 13 blocks. Each block started with five “top-up” adaptation trials. These top-up trials are identical to those in the adaptation phase to help participants recap the adaptation, counteracting potential decay of the recalibration effect during the testing phase. For the subsequent testing trials, a random delay interval of 0, 50, 100, or 150 ms was introduced. Each delayed interval was tested equally within each block (three repetitions in Experiment 1 and four repetitions in Experiment 2). No accuracy feedback was provided for the reproductions for these testing trials.

Accuracy feedback was provided *only* during the Adaptation phase or ‘top-up’ adaptation trials in the testing phase. To encourage participants to rely on internal prior rather than the duration of the contingent sensory event, they received accuracy feedback in text for 400 ms after each reproduction during the adaptation phase. The feedback indicated whether their reproduction was “too short,” “good,” or “too long” based on their relative error. The “good” window was defined as ±10% Weber fraction around 800 ms, following the convention of previous studies (e.g., Ganzenmüller et al., 2012). Specifically, reproduced motor durations between 720 to 880 ms were considered as “good”. Reproduced duration outside this window were marked as either “too short” or “too long”.

### 2.5. Data Analysis

To equalize the number of trials across participants, we used only the last 60 trials from the adaptation phase and the testing trials for further data analysis. We then calculated the mean reproduced durations separately for each session (delayed / synchronized), phase (adaptation / testing), sensory feedback delay, and individual participant. Statistical analyses were conducted in R (R Core Team, 2022). To evaluate the effects of session, phase, and delay on reproduction times, we performed repeated-measures analyses of variance (ANOVAs). We then used linear regression to model the integration of visual and tactile sensory feedback with motor reproduction and to quantify the relationship between introduced delay and reproduced duration. Post-hoc pairwise comparisons were conducted using t-tests with Bonferroni correction where appropriate.

## 3. Result

The following results section presents data from both Experiments 1 (visual feedback) and 2 (tactile feedback) together, as they employed the same paradigm. Findings common to both modalities are reported first, followed by a direct comparison of the effects of sensory modality. This section is structured as follows: we first analyze the adaptation and testing phases separately, examining the effects of delay and modality. We then directly compare these phases to quantify the effect of removing accuracy feedback. Finally, we model the underlying sensorimotor integration processes that gave rise to the observed temporal overestimation.

### 3.1. Adaptation Phase: Duration Reproduction with Accuracy Feedback

First, we examined if accuracy feedback during the adaptation phase could train participants to ignore the 150-ms delay. We compared the mean reproduction durations between the synchronized and delayed sessions and the tactile and visual modalities. The mean reproduction durations (±SEs) during the adaptation phase were 829±17.7 ms for Visual/Delayed, 843±6.3 ms for Tactile/Delayed, 782±12.1 ms for Visual/Synchronized, and 788±8.7 ms for Tactile/Synchronized conditions (Figure 3A). A mixed ANOVA with the within-subject factor Delay (Delayed, Synchronized) and the between-subject factor of Modality (Visual, Tactile) revealed a significant main effect of Delay (*F*(1, 36) = 26.16, *p* < .001, 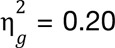). Neither the main effect of Modality (*F*(1, 36) = 0.52, *p* = .475, 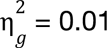) nor the interaction between Delay and Modality (*F*(1, 36) = 0.17, *p* = .678, 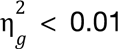) were significant. These findings suggest that, despite accuracy feedback for each trial, the delay still lengthened the reproduction action by 46±17.4 ms for the visual modality and 54±9.1 ms for the tactile modality (Figure 3C). Assuming the general bias was the same for the delayed and synchronized session, based on Equation (1), this lengthening of the motor response corresponded to the integration of about one-third of the 150 ms delay into the motor time (Visual: 0.307±0.12; Tactile: 0.362±0.06). On the other hand, accuracy feedback during the delayed adaptation informed action to shorten the contingent sensory event by two-thirds of the delayed time. Interestingly, there was no significant difference between the visual and tactile modality (*t*(18) = −0.48, *p* = .638, *Cohen’s d* = −0.14, *BF* = 0.338), suggesting that accuracy feedback might dominate sensorimotor recalibration during the adaptation phase, making the influence of modality precision negligible.

**Figure 3.**
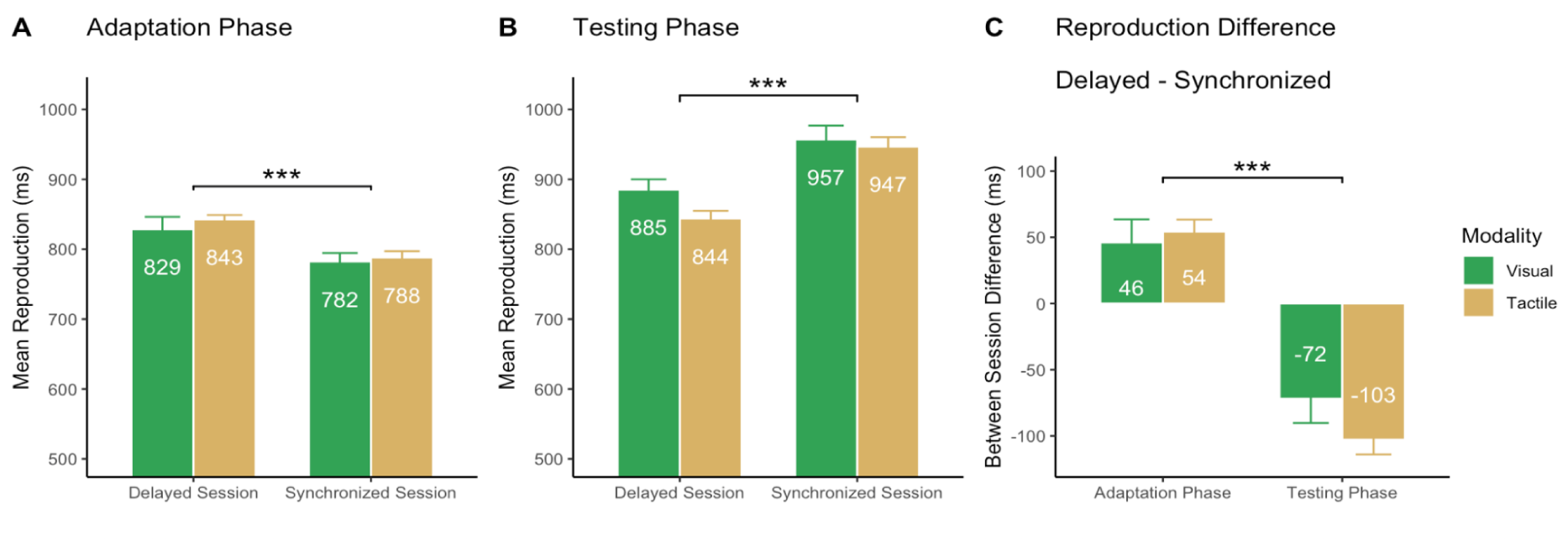
Mean reproduction durations from the adaptation phases (A) and testing phase (B), separated by Session (Delayed / Synchronized) and Modality (Visual: green; Tactile: brown). Panel (C) shows the mean reproduction duration differences between sessions (with Synchronized - Delayed), separated for each phase (Adaptation/Testing) and modality. Asterisks mark the significant between-session difference (***: *p* < 0.001).

### 3.2. Testing Phase: Duration Reproduction without accuracy feedback

After the adaptation phase, the delay between reproduction and contingent sensory feedback varied between 0 ms and 150 ms, critically, with no accuracy feedback. The mean reproduction durations (±SEs) were 885±14.9 ms for the Visual/Delayed, 844±11.0 ms for Tactile/Delayed, 957±19.9 ms for Visual/Synchronized, and 947±13.5 ms for Tactile/ Synchronized, respectively (Figure. 3B). A mixed ANOVA with the same factors as those used in the adaptation phase revealed a significant main effect of Delay (*F*(1, 36) = 66.75, *p* < .001, 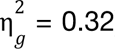). However, neither the main effect of Modality (*F*(1, 36) = 1.92, *p* = .175, 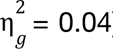) nor the interaction effect between the Delay and Modality (*F*(1, 36) = 2.10, *p* = .156, 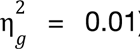) were significant. Interestingly, in contrast to the adaptation phase, the synchronized session yielded longer reproduced duration than the delayed session. Next, we further analyzed across phase differences.

### 3.3. Adaptation versus Testing (or with versus without Accuracy Feedback)

A three-way mixed ANOVA with between-participant factor Modality (Visual, Tactile) and within-participant factors Accuracy Feedback (With, Without) and Adaptation (Synchronized, Delayed) on the mean duration reproduction revealed significant main effects of Accuracy Feedback, *F*(1,36) = 168.87, *p* < .001, 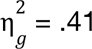, and Adaptation, *F*(1,36) = 15.32, *p* <.001, 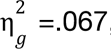, while there was no significant difference between modalities, *F*(1,36) = 0.925, *p* = .343, 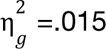. On average, the absence of accuracy feedback during the testing phase lengthened the mean reproduction by 99±8.27 ms compared to the adaptation phase with accuracy feedback. Additionally, the delayed adaptation phase increased the mean reproduction by 50±9.71 ms compared to the synchronized adaptation.

Notably, the interaction between Accuracy Feedback and Adaptation was significant, *F*(1,36)=149.05, *p* < .001, 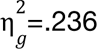. This interaction reflects the opposite patterns shown in Figures. 3A and 3B. To better capture the interaction and opposite effect, we replotted the difference between the two sessions (Delayed - Synchronized, Figure. 3C). Compared to the synchronized session, the reproduction in the delayed session was lengthened by about one-third of the 150-ms delayed duration in the adaptation phase and shortened by two-third of the delayed duration in the testing phase. These results suggest an adaptation toward a shortened sensory feedback timing in the delayed session, resulting in shorter reproductions for the delayed session in the testing phase when both synchronized and delayed sessions experienced the same delay variations.

There was also a significant three-way interaction between Modality, Accuracy Feedback, and Adaptation, *F*(1,36) = 7.7, *p* = .0087, 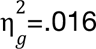. This three-way interaction was likely due to modality modulation in the testing phase but not in the adaptation phase. We will investigate this in detail in the next “sensorimotor integration” session.

### 3.4. Overestimation in the Absence of Accuracy Feedback

The cross-phase comparison revealed that the absence of accuracy feedback increased the mean reproduction duration. However, this initial analysis of the overall mean did not account for the different delay manipulations between the adaptation phase (which used a fixed delay of 150 ms) and the testing phase (which used varied delays of 0, 50, 100, and 150 ms). To enable a fair comparison, we selected only the trials from the testing phase that matched the adaptation conditions: specifically, we compared the 150-ms delay trials from the delayed session with the 0-ms delay trials from the synchronized session. To better visualize the lengthening effect in the absence of accuracy feedback, we subtract the reproduction duration in the adaptation phase from the testing phase. Figure 4 depicts the lengthening effect in the testing phase compared to the adaptation phase.

**Figure 4.**
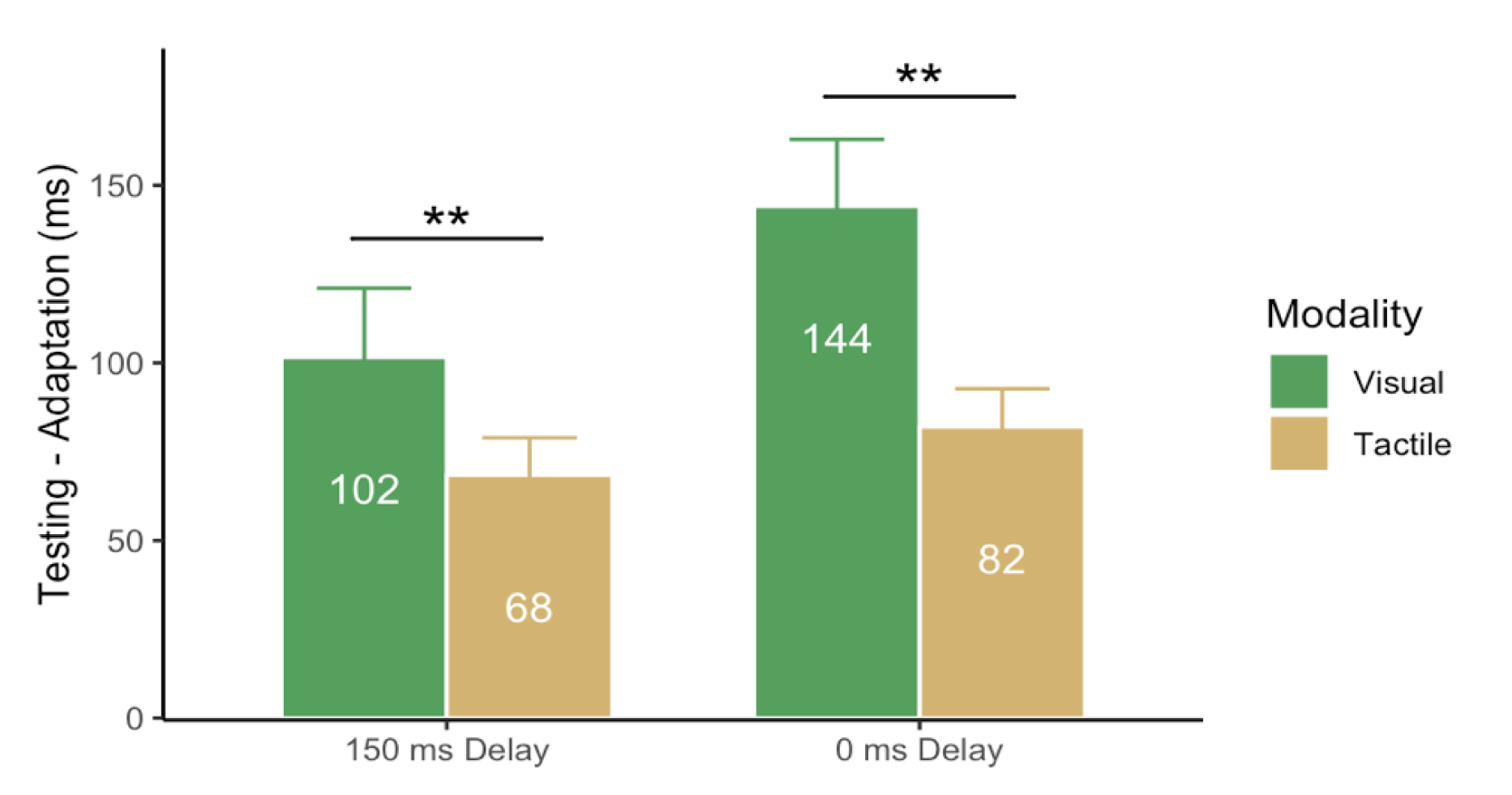
The mean lengthening effect (Testing - Adaptation) for the delayed (150-ms trials only) and synchronized (0-ms trials only) sessions, separated for the visual (green) and tactile (brown) modality. The asterisks mark the significant between-session difference (**: p < 0.01).

A two-way (Session: Delayed, Synchronized) × Modality (Visual, Tactile) ANOVA analysis revealed a significant main effect of Modality, *F*(1, 36) = 9.94, *p* = .003, 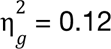. However, neither between session main effect (*F*(1, 36) = 3.14, *p* = .085, 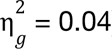) nor the interaction effect (*F*(1, 36) = 0.84, *p* = .366, 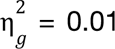) were significant. Importantly, the lengthening effects were all significantly positive (*ts* > 5, *ps* < .001). These results suggest that without accuracy feedback to recalibrate the motor timing in the testing phase, the motor reproduction shifted back to its original state, exhibiting a general overestimation consistent with previous findings (Chen & Shi, 2023; Shi, Ganzenmüller, et al., 2013). Interestingly, this reverting was less prominent for tactile than visual reproduction, indicating reproducing tactile durations were relatively more robust in the absence of accuracy feedback.

### 3.5. Sensorimotor Integration in the Absence of Accuracy Feedback

To quantitatively assess differential weights of action-induced visual versus tactile timing in the absence of accuracy feedback, we further conducted linear mixed regression analyses. As stated in Introduction, the sensorimotor integration with delay manipulation can be expressed as Equation (3): *M* = *D*_*p*_ + *vD*, where D_p_ is the perceived duration, and D is the delay. Figure. 5 shows the linear trends with the delay manipulations in the testing phase, separated for the visual (green) and tactile (brown) sensory feedback, as well as the delayed and synchronized sessions. The trends were consistent within each modality, but the trend with tactile stimuli was steeper than the visual stimuli.

**Figure 5.**
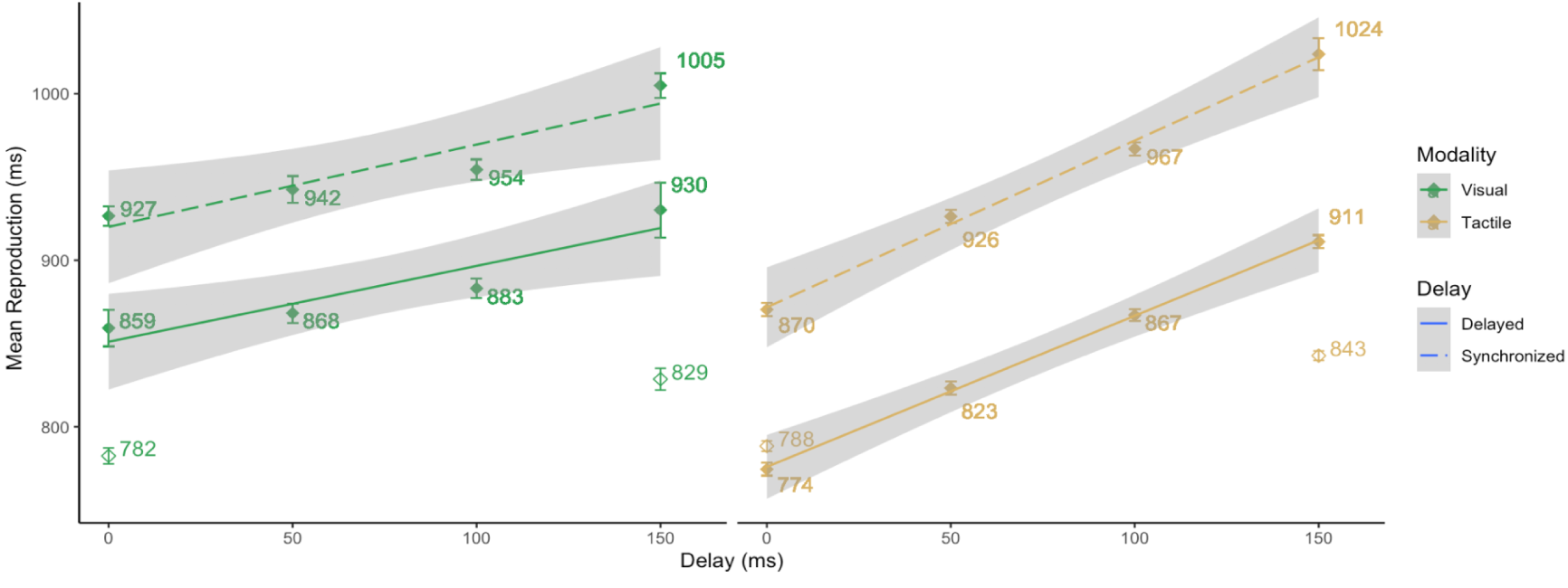
The mean reproduction durations (solid points) in the testing phase as a linear function of delay, separated for the delayed (solid lines) and synchronized (dashed lines) sessions, and Visual (left panel) and Tactile (right panel). The un-filled points indicate the mean reproduction duration in the adaptation phase. The mixed linear regression analysis revealed the slope was significantly steeper for the tactile (0.95) than visual (0.47) sensory feedback.

Based on these observations, we conducted linear mixed regression, assuming the delay adaptation only changed the internal prior, while the linear trend was determined by the weight of the sensory feedback in sensorimotor integration. That is, we conducted the following linear mixed-effects model within each modality:

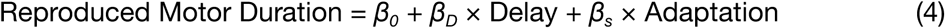

where Adaptation is a dummy variable (0 - delayed, 1 - synchronized) for the two sessions.

The estimated coefficients and linear models are as follow:

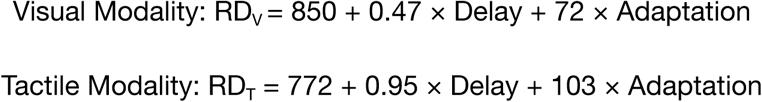

Both coefficients of Delay were significant, (Visual: *t*(18) = 8.44, *p* < .001, Tactile: *t*(18) = 17.87, *p* < .001). Compared across modalities, the slope was significantly higher with tactile than visual durations (*t*(18) = 6.95, *p* < .001, *Cohen’s d* = 2.05, *BF* = 41084.52), indicating that duration reproduction relied more heavily on the contingent sensory timing with tactile than with visual stimulation in the absence of accuracy feedback.

Comparing the delayed vs. synchronized sessions, delayed adaptation significantly shortened the reproduction in the testing phases in both modalities (Visual: *t*(18) = 3.93, *p* < .001, Tactile: *t*(18) = 9.27, *p* < .001), consistent with the mean reproduction analysis. The overall shortening was not significantly different between the tactile and the visual modality (*t*(18) = 1.550, *p* = 0.15, *Cohen’s d* = 0.47, *BF* = 0.713). These results indicated that both delayed duration and adaptation had significant impact on the reproduction duration. Delay adaptation shortened the internal prior, which remained effective during the testing phase.

Taken together, these results revealed a consistent pattern of sensorimotor integration in both visual and tactile reproduction. The reproduction duration increased linearly as the given delay duration increased in the testing phase, independent of the delay adaptation effect, which was also prominent in both experiments.

## 4. Discussion

The present study aimed to investigate the impact of accuracy feedback on delay adaptation and the weighting of motor versus sensory feedback timing in visual and tactile duration reproduction. During the adaptation phase, one session introduced a 150 ms sensory feedback delay, while the other used synchronized sensory feedback. Critically, accuracy feedback was provided during the adaptation phase but not in the testing phase. In the testing phase, the delay of the contingent sensory feedback varied from 0 to 150 ms, allowing us to use linear approximation to estimate the weights of sensorimotor integration in duration reproduction. Our findings indicated that accuracy feedback recalibrated the motor action, although one-third of the delay was still integrated into motor reproduction. Importantly, in the testing phase without accuracy feedback, the recalibrated internal prior from the delayed adaptation session continued to influence performance: participants who experienced delayed feedback during adaptation produced systematically shorter reproductions in the testing phase compared to those who experienced synchronized feedback during adaptation (by 72 ms for visual and 103 ms for tactile modality). This persistent effect demonstrates that the adaptation-induced changes to internal prior representations were maintained even when accuracy feedback was removed. Additionally, sensorimotor integration revealed that duration reproduction without accuracy feedback relied more heavily on the contingent sensory tactile event than the visual event. The results demonstrate a consistent delay adaptation effect across both modalities, but the extent of adaptation and sensorimotor integration differed between visual and tactile feedback.

### 4.1. Influences of accuracy feedback

During the adaptation, participants received accuracy feedback at the end of each trial to encourage reliance on their motor timing rather than sensory feedback timing. Previous research showed that in the absence of accuracy feedback, the total duration of motor reproduction (from button press to release) was immediately lengthened to match the onset delay from the very first trial when the delay was introduced (Ganzenmüller et al., 2012). This immediate lengthening suggests participants relied primarily on matching the durations of the probe stimulus and action-induced sensory feedback (within-modality sensory comparison) rather than on internal prior. Our study demonstrates that accuracy feedback was a critical factor for altering this strategy and recalibrating internal prior and thus motor timing, although participants could not completely discount the delay even with explicit accuracy feedback. Specifically, the reproduced duration was notably longer (with a session difference of 47 ms and 55 ms for the visual and tactile feedback, respectively) in the delayed feedback sessions compared to the synchronized ones. This indicates that approximately one-third of the delay (150 ms) was still integrated into motor reproduction, while accuracy feedback successfully encouraged participants to compensate the remaining two-thirds (approximately 100 ms). This partial compensation demonstrates that sensory feedback was still factored into the sensorimotor reproduction, consistent with early research (Shi, Ganzenmüller, et al., 2013).

A significant distinction between the adaptation and testing phase was the lack of the accuracy feedback during the testing trials. Consequently, participants had to rely on their own pre-existing prior, recalibration during the adaptation, and balance the integration weights of the motor timing and contingent sensory feedback timing. Without the guiding accuracy feedback, as evidenced by the linear mixed models, the session difference (synchronized - delayed) remained to be large: 72 ms for the visual and 103 ms for the tactile modality. This suggests the recalibration remained ineffective in the testing phase when the accuracy feedback is missing .

### 4.2. Modality-specific sensorimotor integration

When comparing visual and tactile sensory feedback, our results revealed modality-specific integration weights that reflect differential reliance on sensory feedback timing versus motor timing. This is evident from the linear slopes derived from the linear mixed model (Equation 3: *M = D_p_ + vD*), with tactile feedback showing a slope of 0.95 compared to the visual feedback’s slope of 0.47. According to the MLE framework and consistent with previous sensorimotor integration research (Ernst & Banks, 2002; Körding & Wolpert, 2004; Shi, Ganzenmüller, et al., 2013), these slopes directly quantify the weight assigned to sensory feedback (v) in the integration process. Our tactile feedback slope of 0.95 demonstrates that participants relied almost entirely on tactile sensory feedback timing, assigning only minimal weight (5%) to motor timing. In contrast, the visual feedback slope of 0.47 indicates more balanced integration, with approximately equal weighting of motor timing (53%) and visual sensory feedback timing (47%).

These modality-specific integration weights align with the broader literature on multisensory reliability (Welch & Warren, 1980): more reliable sensory information receives greater weight in determining the integrated percept. The near-complete reliance on tactile feedback suggests that tactile temporal information was perceived as highly reliable for this duration reproduction task, consistent with findings that tactile information dominates over visual information in temporal tasks (Bresciani et al., 2006). The more balanced weighting for visual feedback suggests that visual temporal information was perceived as less reliable, leading the sensorimotor system to integrate it more equally with motor timing information.

One might argue that this heavy reliance on tactile stimulation is due to close proximity setup, as the tactile feedback was closer to the response finger compared to the visual feedback. The spatial proximity has been shown to be a critical factor for multisensory integration (Chen et al., 2018; Koelewijn et al., 2010). On-body touch stimulation is often associated with general defensive action (Yeomans et al., 2002), so we expect that stimulation on other body parts would yield similar findings as stimulation on the response finger. Additionally, on-body touch stimulation often induces higher valence than the neutral visual stimulation, which might contribute to the higher weight of the tactile feedback compared to visual feedback. However, the present findings did not differentiate between stimulation of the same versus different body parts. Future research should clarify this distinction.

### 4.3. Shifts of the prior with varied delayed feedback

In both experiments we included a control session with synchronized sensory feedback during the adaptation phase. This session enabled us to examine influences of accuracy feedback (present during the adaptation but absent during the testing) and varied delayed feedback on prior formation.

With the accuracy feedback, both visual and tactile duration reproduction were relatively close to the target duration of 800 ms (visual at 782 ms, tactile at 788 ms). However, in the absence of the accuracy feedback, reproduction with synchronized sensory feedback in the testing phase was lengthened to 927 ms for visual feedback and 870 ms for tactile feedback. Two potential contributing factors may account for this lengthening effect. First, the overestimation may arise from divided attention during reproduction. As demonstrated in a recent study (Chen & Shi, 2023) that compared the presence and absence of accuracy feedback, the lack of accuracy feedback lengthened reproduction by about 13.5% in both subsecond and supra-second reproductions. The second contributing factor is the perturbation of various delays. Since the delays we equally administered ranged from 0 to 150 ms, the averaged delay was about 75 ms in the testing phase. This means, the sensory feedback was shortened by an average of 75 ms compared to the adaptation phase. Given that the prior developed in the adaptation (the synchronized session) was based on full synchronized feedback, introducing such delays in the testing phase lengthened the reproduced duration to equate the perceived duration. As a result, the mean reproduction in the testing phase expanded nearly 150 ms for both modalities, resulting in 957 ms for visual and 947 ms for tactile reproductions. It is noteworthy that both modalities had nearly identical mean reproductions, suggesting that observers might perceive and encode durations similarly across visual and tactile modalities. Nevertheless, the impact of the sensory feedback and delay varied, evidenced by the differing integration weights we observed. Dissecting these two components to understand their individual influences, however, requires future research.

It is important to note that the lengthening trends in the synchronized sessions mirrored those in the delayed feedback sessions, given that we didn’t observe any interactive effects. The overestimation in terms of ratio was larger in the synchronized feedback session (Visual: 18.5%, Tactile: 10.4%) and smaller in the delayed feedback session (Visual: 12.2%, Tactile: 8.1%). This implies that the acquisition of the prior (with or without delay) during the adaptation phase acts additively to the overestimation brought about by the absence of the accuracy feedback and introduced delays. In other words, adaptation of delay sensory feedback influenced the prior (that is, how the standard duration is perceived), while the trial-to-trial variation in delay primarily impacted the sensorimotor integration. Without the accuracy feedback, those trial-to-trial sensorimotor integrations (with the delay) progressively update the internal prior, resulting in an overestimation that we observed.

### 4.4. Limitation and future directions

While the current study focuses on modality-specific delay adaptation, several limitations warrant consideration. First, the tactile stimulus was placed on the response finger for spatial alignment, which may have enhanced tactile salience and influenced its perceptual weighting relative to visual feedback. Future research should systematically vary stimulus location (action vs. non-action finger, hand vs. arm) to isolate the effect of spatial proximity from modality-specific reliability. Second, the current study employed only unimodal conditions; cross-modal visual-tactile stimuli would provide valuable insights into how the brain resolves conflicts between modalities during temporal recalibration. Third, we used a single standard duration (800 ms). While well-established in the literature, exploring a wider range of durations (both sub-second and supra-second) would clarify the generalizability of delay adaptation effects across different timing mechanisms. Finally, the divided attention demands of monitoring sensory feedback while reproducing duration may have contributed to overestimation effects observed without accuracy feedback. Future work should dissociate the contributions of attentional load from pure sensorimotor recalibration.

### 4.5. Conclusion

In conclusion, this study examined the effects of delayed sensory feedback on visual and tactile duration reproduction. We observed consistent adaptation effects with accuracy feedback for both modalities, but different afftereffects without accuracy feedback. In the absence of accuracy feedback, tactile reproduction relied more on sensory feedback and ignored the delay between action and sensory feedback, resulting in greater lengthening of reproduction compared to the visual modality. Additionally, overestimation occurred in the absence of accuracy feedback and with the presence of delay. Overall, our findings suggest that temporal delay adaptation is influenced by accuracy feedback and sensorimotor integration. The integration was governed by sensory reliability and weighted differently across modalities, with a higher weight on the tactile feedback than visual feedback.

## Data Availability Statement

None of the experiments was pre-registered. The data and materials for experiments are available at: https://github.com/msenselab/Aftereffect_Delay_Adaptation.

